# Global Plant Extinction Risk Assessment Inform Novel Biodiversity Hotspots

**DOI:** 10.1101/2021.10.08.463027

**Authors:** Thomas Haevermans, Jessica Tressou, Joon Kwon, Roseli Pellens, Anne Dubéarnès, Simon Veron, Liliane Bel, Stéphane Dervaux, Juliette Dibie-Barthelemy, Myriam Gaudeul, Rafaël Govaerts, Gwenaël Le Bras, Serge Muller, Germinal Rouhan, Corinne Sarthou, Lydie Soler

## Abstract

Curbing biodiversity loss and its impact on ecosystem services, resilience and Nature’s Contributions to People is one of the main challenges of our generation (IPBES, 2019b, 2019a; Secretariat of the United Nations Convention on Biological Diversity, 2020). A global baseline assessment of the threat status of all of biodiversity is crucial to monitor the progress of conservation policies worldwide (Mace & al., 2000; Secretariat of the United Nations Convention on Biological Diversity, 2021) and target priority areas for conservation (Walker & al., 2021). However, the magnitude of the task seems insurmountable, as even listing the organisms already known to science is a challenge (Nic Lughadha & al., 2016; Borsch & al., 2020; Govaerts & al., 2021). A new approach is needed to overcome this stumbling block and scale-up the assessment of extinction risk. **Here we show** that analyses of natural history mega-datasets using artificial intelligence allows us to predict a baseline conservation status for all vascular plants and identify target areas for conservation corresponding to hotspots optimally capturing different aspects of biodiversity. We illustrate the strong potential of AI-based methods to reliably predict extinction risk on a global scale. Our approach not only retrieved recognized biodiversity hotspots but identified new areas that may guide future global conservation action (Myers & al., 2000; Brooks & al., 2006). To further work in this area and guide the targets of the post-2020 biodiversity framework (Díaz & al., 2020a; Secretariat of the United Nations Convention on Biological Diversity, 2020; Mair & al., 2021), it will be necessary to accelerate the acquisition of fundamental data and allow inclusion of social and economic factors (Possingham & Wilson, 2005).

## Introduction

Assessing the threat status of all known species is a stated objective to guide conservation actions across the world (UNEP (United Nations Environment Programme) & CBD (Convention on Biological Diversity), 2010) and enable the sustainable management of life on Earth (Newbold & al., 2016). Recent assessments indicate that the window to reverse the current trends of biodiversity loss, in a serious commitment to intergenerational justice, is narrowing fast (IPBES, 2019b, 2019a; Secretariat of the United Nations Convention on Biological Diversity, 2021). These assessments emphasize that the importance of rapidly obtaining high quality data for guiding global conservation policies. As a result, the United Nations Convention on Biological Diversity (CBD) is now working on a new generation of targets, goals and indicators aimed at curbing species extinction and reversing biodiversity loss (Díaz & al., 2020a; Secretariat of the United Nations Convention on Biological Diversity, 2021).

Extinction risk and biodiversity distribution data form the baseline for tracking our progress within the CBD’s post-2020 biodiversity framework. The Global Strategy for Plant Conservation (GSPC), included within the CBD in 2002, is a central project for documenting and conserving plant diversity. Through its actions, massive datasets of occurrences are now available for assessment and analysis. Unfortunately, due to the sheer scale of the task, the evaluation of plant conservation statuses by the International Union for the Conservation of Nature (IUCN) and other national and international bodies is lagging behind, and a global threat status is still lacking for the vast majority of organisms. Indeed, despite the recognized importance of species extinction risk evaluation, the assessment data available is still far from sufficient to enable the establishment of clear goals and track progress within the post-2020 framework, which aims to promote science-based targets for biodiversity conservation (Mair & al., 2021). Currently, only 138,374 species have a global status on the IUCN Red List (IUCN, 2021), representing only *ca*. 7% of the estimated number of described species, with a strong bias towards certain geographical regions or flagship groups, for which datasets are the most complete, such as vertebrates and trees (Bachman & al., 2019), and whose permanent loss is perceived as having an impact on humanity (IPBES, 2019a). This figure includes *ca*. 56,000 plant species, which represents less than 15% of the total number of plants known to science. So far, most global-scale conservation assessments have focused on vertebrates (Ceballos & al., 2020; Robuchon & al., 2021), even though global conservation planning should be informed by more species-rich groups such as invertebrates (Mace & al., 2000) and vascular plants, which are of outstanding and economic importance for human well-being (Molina-Venegas & al., 2021). The overwhelming number of species with insufficient data along with the difficulties in reassessing species in the field hamper any rapid progress and make it currently impossible to assign a formal threat status to an important fraction of existing species, which will, at best, remain classified as Data Deficient (Mair & al., 2021).

Machine learning methods (AI) can find predictive patterns that humans would not have been capable of spotting or identifying as relevant within huge and highly dimensional datasets (Walker & al., 2021). Natural history accessions and their associated information (locality, time of collection), as well as the taxonomic history of each species represent a as of yet unexplored source of information and may be potential predictors of frequency, distribution range and period during which a species was observed (Albani Rocchetti & al., 2021). The recent digitization of major global natural history collections (Le Bras & al., 2017) has given access to a wealth of information (Paton & al., 2020) and has contributed to authoritative expert-based lists of known plant life (Le Bras & al., 2017; Borsch & al., 2020; Govaerts & al., 2021). We used machine learning methods to explore an innovative, objective and reproducible method for assigning a threat status to all known vascular plant (VP) species in one go using massive datasets (Pelletier & al., 2018; Nic Lughadha & al., 2020). Our results provide the first global assessment of VP conservation status and define a new set of science-based Plant Conservation Hotspots (PCH2.5 and PCH5, see Methods), taking into account three commonly used biodiversity facets (Myers & al., 2000; Forest & al., 2007).

## Results

Performance (test scores) of the AI algorithm in Extinction Risk (ER) prediction for *n* = 345,697 species (**Supplementary Table 4**) was 82.7%, 70%, and 53% for the two, three and five classes, respectively. The importance of each of the 535 features in the prediction models is given in **Supplementary Table 2**. Results were distributed as follows and were converted into threatened (Þ) / non-threatened (NÞ) status using the thresholds defined by the IUCN (*i*.*e*. LC.NT = NÞ and VU.EN.CR = Þ, see Methods): two classes (41% LC.NT, 59% VU.EN.CR, corresponding to 41% NÞ / 59% Þ), three classes (30% LC, 20% NT, 50% VU.EN.CR, corresponding to 50% NÞ / 50% Þ), and five classes (26% LC, 20% NT, 8% VU, 9% EN, 37% CR, corresponding to 46% NÞ / 54% Þ). When considering specific plant groups such as trees (here, ‘Phanerophytes’), we find that our results are congruent with the most recent global estimates (Botanic Gardens Conservation International, 2021). Indeed, one third of trees are threatened worldwide (44% when including the ‘possibly threatened’ category), and up to two thirds are threatened in Madagascar. Those figures match our own results (59% of trees are threatened in Madagascar, 41% globally (*n* = 89,151)). The percentage of species assigned to the DD category in the training dataset that were predicted to be threatened ranged between 65– 73% (2 classes: all 59% Þ / DD 73% Þ, 3 mod.: all 50% Þ / DD 65% Þ, and 5 mod.: all 54% Þ / DD 71% Þ), a proportion that is in between the “best” and the “high” estimates from the IUCN (IUCN, 2021) (*i*.*e*. best = DD species are predicted to be threatened in the same proportion as data sufficient species; high = DD species are always predicted to be threatened). The top 2.5% and 5% hotspot areas (using botanical countries normalized for size, see Methods) for each facet (current: SPR, PD, END; threatened: ÞSPR, ÞPD, ÞEND and/or expected losses 50 years from now: ℓSPR, ℓPD, ℓEND) are listed in **Extended Data Table 4**. Analyses of the correlation between facets revealed two main groups (**Figure 1**, rankings given in **Supplementary Table 3**).

**Figure 1.**
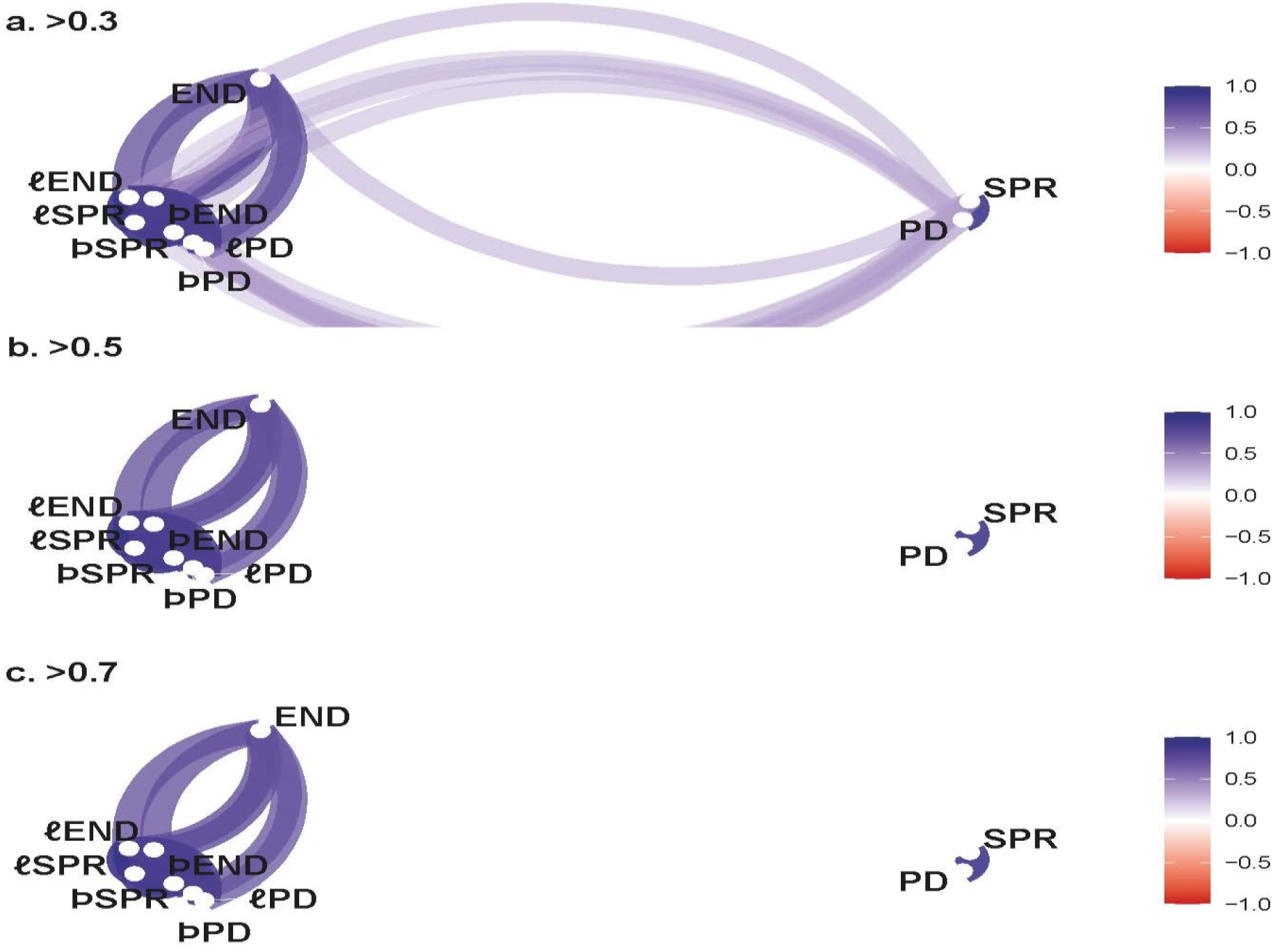
Correlogram (R package *Corrr*) of all biodiversity facets obtained using Kendall’s *τ*. (**a**) weak correlation (>0.3). (**b**) moderate correlation (>0.5). (**c**) high correlation (>0.7). The correlation between variables is indicated by clustering and color intensity. Clustering: *(i)* SPR and PD are strongly correlated (0.89) while only weakly correlated (0.39– 0.40) with endemicity (END), *(ii)* the various measures of threat (Þ and ℓ) are all strongly correlated (0.78–0.93) and are correlated with END (0.71–0.75) but only weakly with SPR or PD (0.32–0.38) (**Extended Data Fig. 4**).

Figure 1 illustrates the existence of two different types of areas, which is confirmed by the PCA rankings of botanical countries (**Supplementary Table 3**) following those biodiversity facets clusters. The Andes were confirmed to be an important PD+SPR hotspot (**Extended Data Fig. 5a**), whereas Old World insular areas were found to be crucial threatened diversity and endemicity hotspots (**Extended Data Fig. 5b**). The SPR and PD hotspots, see **Extended Data Table 4**, strongly overlapped (only differing by one country) and were mostly located in continental areas in the Neotropics, with Colombia having the highest SPR/PD (**Extended Fig. 1a and 1c; Supplementary Table 3**). The END hostpots were mostly insular and paleotropical areas, with Madagascar (MDG) having the highest END in the world (0.825) (**Extended Data Fig. 1b, Supplementary Table 3**). Hotspots of threatened diversity (ÞSPR, ÞPD, and ÞEND) and with expected losses of diversity (ℓPD, ℓPD and ℓEND) overlapped, albeit not completely, with END hotspots, being mostly paleotropical with almost half occurring on insular territories (**Extended Data Figs. 2 and 3)**.

Priority Conservation Hotspots (PCH) identified from PCA ordination with 2.5% and 5% thresholds (PCH2.5 and PCH5) capture most of the contribution of each facet to global plant biodiversity (**Fig. 2**). PCH capture a similar proportion of each facet than the top 2.5% and 5% area identified for each facet (**Extended Data Table 2**), but with a notable minimal taxonomic overlap as indicated in **Extended Data Table 3**. PCH thus capture top endemics from SPR and PD hotspots, as well as top endemics from END hotspots (by definition containing non-overlapping taxa strictly restricted to distinct areas).

**Figure 2.**
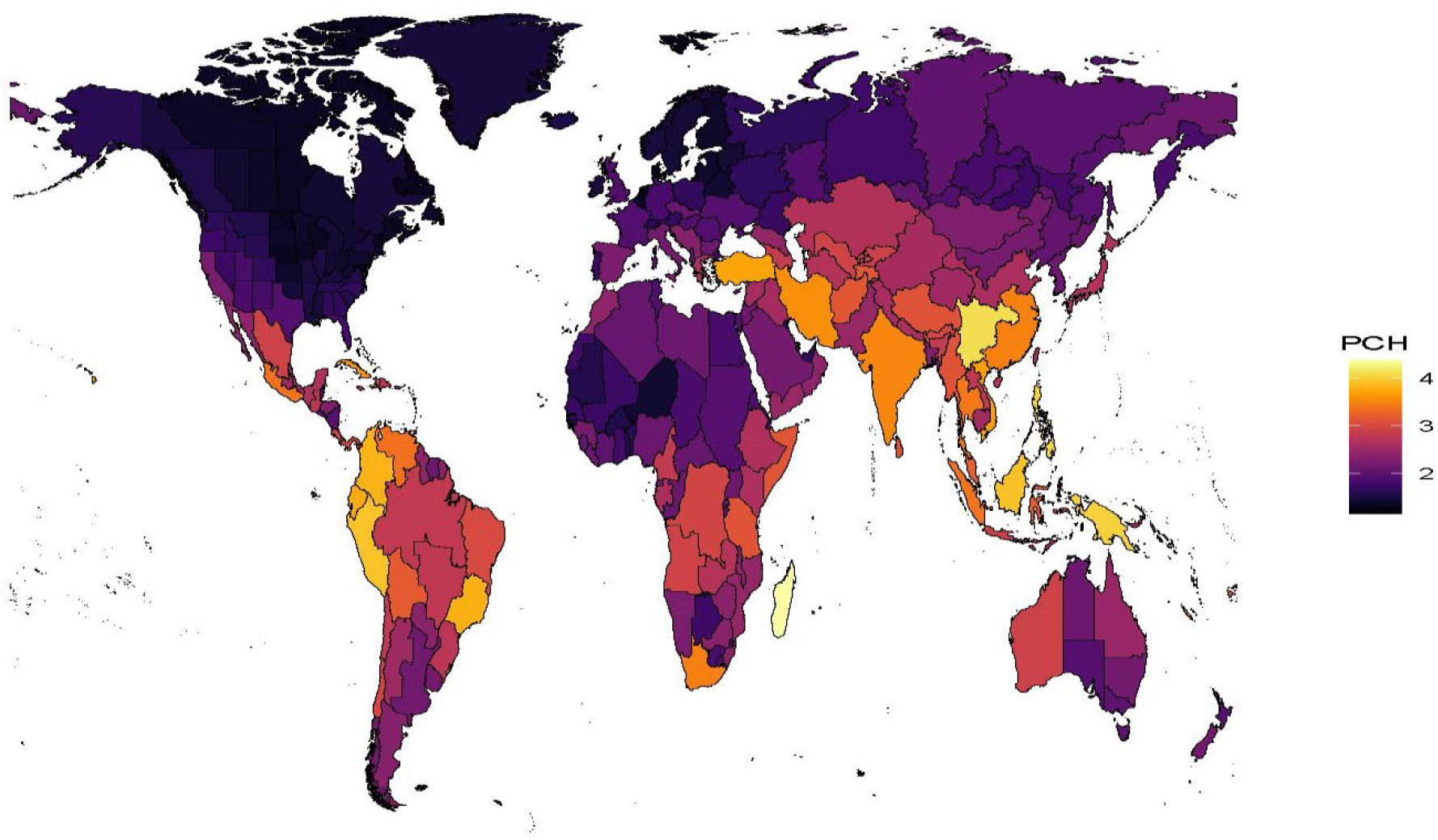
Map of Priority Conservation Hotspots. PCH ranked according to values of axis 1 of the PCA analysis of all variables (see Methods), taking into account both the hyperdiversity and irreplaceability of the species harbored by botanical countries.

Our results show that Madagascar, China South-Central, the Philippines, New Guinea, Peru, Borneo, Colombia, Brazil Southeast and Ecuador, identified as our PCH2.5, are of extreme importance for their global contribution to plant biodiversity (**Fig. 2**, see complete rankings in **Supplementary Table 3**). PCH5 also included Turkey, Hawai‘i, Vietnam, Cuba, Iran, India, China Southeast, the Cape Provinces and Thailand. The total area covered by PCH2.5 represents *ca*. 5% of emerged areas (*n* = 114,577 distinct taxa, *i*.*e. ca*. 30% of VP) and PCH5 represents up to 10% (*n* = 164,142 taxa, *i*.*e. ca*. 50% of VP) (**Extended Data Tables 3 and 4**).

Overall, our analyses suggest that PCH5 contain 50% of the world’s species richness, 43% of the plant tree of life and 53% of the world’s highly endemic species. These areas also contain a quarter of threatened species (ÞSPR), almost half of the threatened tree of life (ÞPD) and a third of the threatened endemics (ÞEND). If no conservation measures are taken to reverse current trends, we estimate that 14% of the global SPR, 27% of endemics and 47% of the plant tree of life may be lost during the next 50 years (ℓSPR, ℓPD and ℓEND in **Extended Table 2**).

### AI and the estimation of the proportion of threatened plants worldwide

Predicting extinction risk is a challenge that has been tackled recently (Humphreys & al., 2019; Nic Lughadha & al., 2020; Walker & al., 2021), and this endeavor is facilitated when local floras are well known and documented, which is not the case for most regions of the world harboring biodiversity hotspots (Myers & al., 2000). Our study is the most comprehensive threat pre-assessment ever carried out for a large clade (*n* = 345,697 vascular plant (VP) species), representing almost all of VP diversity. The large number of features used here (535) ensures that we can identify the features that may be relevant for performing the prediction task (the importance of each feature is given in **Supplementary Table 2)**. The use of sophisticated AI methods is especially relevant for this kind of projection by efficiently managing raw biodiversity data (Walker & al., 2021). Climate features, in particular those linked to aridity (‘Climate_Desert’, ‘Climate_Consistently_Dry’) were important in two of our prediction schemes. However, the most highly ranked features in the final models were distribution area-related parameters, including several artificial features created by a neural network (**Supplementary Tables 1 and 2)**. Indeed, we expected that some of these artificial features would account for the quantity of occurrence data as well as the geographical range of species, and therefore be correlated to the threat level, since the IUCN Red List criteria are based on such parameters. The IUCN takes into account low population size and the tendency to decline (criteria A, C, D), geographical range (B), or uses a quantitative analysis (Mace & al., 2008) indicating a probability of extinction in the wild (E). However, in practice most of the Red List assessments strongly rely on distribution data due to the difficulty in obtaining reliable measurements for most criteria, resulting in threat pre-assessments for plants being *de facto* based on criterion B (Pelletier & al., 2018; Stévart & al., 2019; Nic Lughadha & al., 2020).

By including geospatial data to account for habitat reduction, previous studies of trees (ter Steege & al., 2015) and birds (Ocampo-Peñuela & al., 2016) suggested that 57% of Amazonian tree species (ter Steege & al., 2015) and nearly 50% of birds (Ocampo-Peñuela & al., 2016) may be threatened, highlighting the fact that earlier methods may have largely underestimated the figures. Other studies of larger datasets (Pelletier & al., 2018; Stévart & al., 2019; Nic Lughadha & al., 2020) of local or global significance provided lower estimates (*e*.*g*. 30% of plant life may be threatened based on spatial data only (Pelletier & al., 2018; Stévart & al., 2019), and 40% based on a MPR (multilevel regressions and post-stratification) method (Nic Lughadha & al., 2020), *i*.*e*. not based on individual assessments). These discrepancies may be explained by the way DD species were accounted for or not, as we showed here that a majority of these species should be considered as threatened (see results). Besides distribution parameters, other features proved to be important in our analyses, in particular taxonomic features (*e*.*g*. number of synonyms, description date, family) and the sampling frequency of a given species (*e*.*g*. number of specimens, sampling date). Our analyses showed that factors other than distribution are relevant for predicting species threat status and should be taken into account when estimating extinction risk.

### Post-2020 targets and priority setting for the conservation of various biodiversity facets

One of the main recommendations of the scientific community for curbing biodiversity loss and reducing the risk of losing many of Nature’s Contributions to People (NCP) (Díaz & al., 2020b) is to set ambitious goals in the post-2020 Global Biodiversity Framework. It is essential to identify hotspots of biodiversity, in its different facets, to define clear targets for reducing species extinctions (Goal A) and conserving ecosystem services and NCP (Goal B). The hotspots we identified (**Extended Data Table 1 and 4**) grouped according to the correlations between facets of biodiversity (**Fig. 1**), illustrating the fact that two main aspects of biodiversity: hyper-diversity (SPR/PD hotspots) and irreplaceability (END/allÞ/allℓ), are not nested (Orme & al., 2005; Díaz & al., 2020b). A high proportion of areas with high endemicity, containing irreplaceable species (Veron & al., 2021), correspond to insular ecosystems, whereas PD and SPR hotspots occur in neotropical continental areas. Surprisingly, the latter harbor a lower proportion of global endemicity (**Extended Data Table 2**). Biodiversity hotspots were first identified by Myers *et al*. from levels of species endemism (with particular focus on vascular plants) and degree of threat (approximated through habitat loss) (Myers & al., 2000). Twenty hotspots are shared (at least partially) between Myers *et al*. and our PCH5 analysis (**Extended Data Table 1**), which is a significant overlap but which nevertheless reveals key differences. The main differences are found in the California Floristic Province, Central Chile, the West African Forests, the Mediterranean Basin, the Caucasus, Southwest Australia, New Zealand and New Caledonia, which are listed in Myers *et al*. (Myers & al., 2000) but are absent from our list. Conversely, we identified Iran, Turkey, India, Southeast China and New Guinea, whereas these areas were not retained by Myers *et al*. or only very partially. Some of these discrepancies might be due to the fact that we used proportions instead of raw counts or to a difference in the way geographical units were defined (we did not separate coastal areas from our botanical countries, unlike Myer *et al*.). Most notable, perhaps, is the absence of New Zealand from the PCH5 list and the inclusion of New Guinea. It is possible that New Guinea would make it on the list if Myers *et al*.’s study was conducted today, due to the catastrophic decline of its Wilderness Areas (Watson & al., 2016) and the richness and irreplaceability of its flora (Cámara-Leret & al., 2020). Some of these metrics are already used in research programs that aim to highlight priority areas for conservation: species extinction risk is one of the criteria for designating Key Biodiversity Areas (KBA) and Important Plant Areas (IPA) (Brooks & al., 2015b; IUCN, 2016; Darbyshire & al., 2017). Thirteen of the PCH5 biodiversity hotspots are in fact already being monitored under the IPA scheme, and numerous KBAs have been identified in all our PCH5 hotspots (BirdLife International, 2021).

Our study fulfill the highly ambitious goals of the CBD for biodiversity and sustainability (Díaz & al., 2020b) by helping optimize the safeguarding of the tree of life and the conservation of a high number of irreplaceable species with globally or locally critically restricted range (Daru & al., 2020), and areas where species diversity is high. Conservation efforts targeting the regions highlighted by PCH5 would likely benefit other branches of the tree of life (twenty of our top 25 PCH overlap with areas that have been shown to be important for the conservation of phylogenetic diversity in mammals (Robuchon & al., 2021)). The individual species threat assessments and the megaphylogeny provided here may also be used at scales other than the one considered here, and may meet other or complementary CBD goals, such as protecting NCPs and ecosystem services (Possingham & Wilson, 2005).

## Data availability

The data sources used to build the features used in this study, the taxonomic and distribution backbone, and the training dataset are detailed in the article; the megaphylogeny along with details of the results are provided in **Supplementary Tables 1–4** and **Supplementary File 1**.

## Code Availability

The R and Python codes used for calculations and analyses are available from the corresponding authors on request.

## Acknowledgements

We thank Hélène Citerne for her help with language editing.

## Author Contributions

TH, JT, JK and RP conceived the study and analyzed the data; SV and TH performed the expected PD loss and PD calculations; TH, JT, JK, RP, AD wrote a first draft of the paper; All authors acquired the data, interpreted the results of the analyses and contributed to writing the final manuscript.

## Additional Information

**Supplementary Information** is available for this paper.

## Extended Data and Figures

**Extended Data Figure 1.**
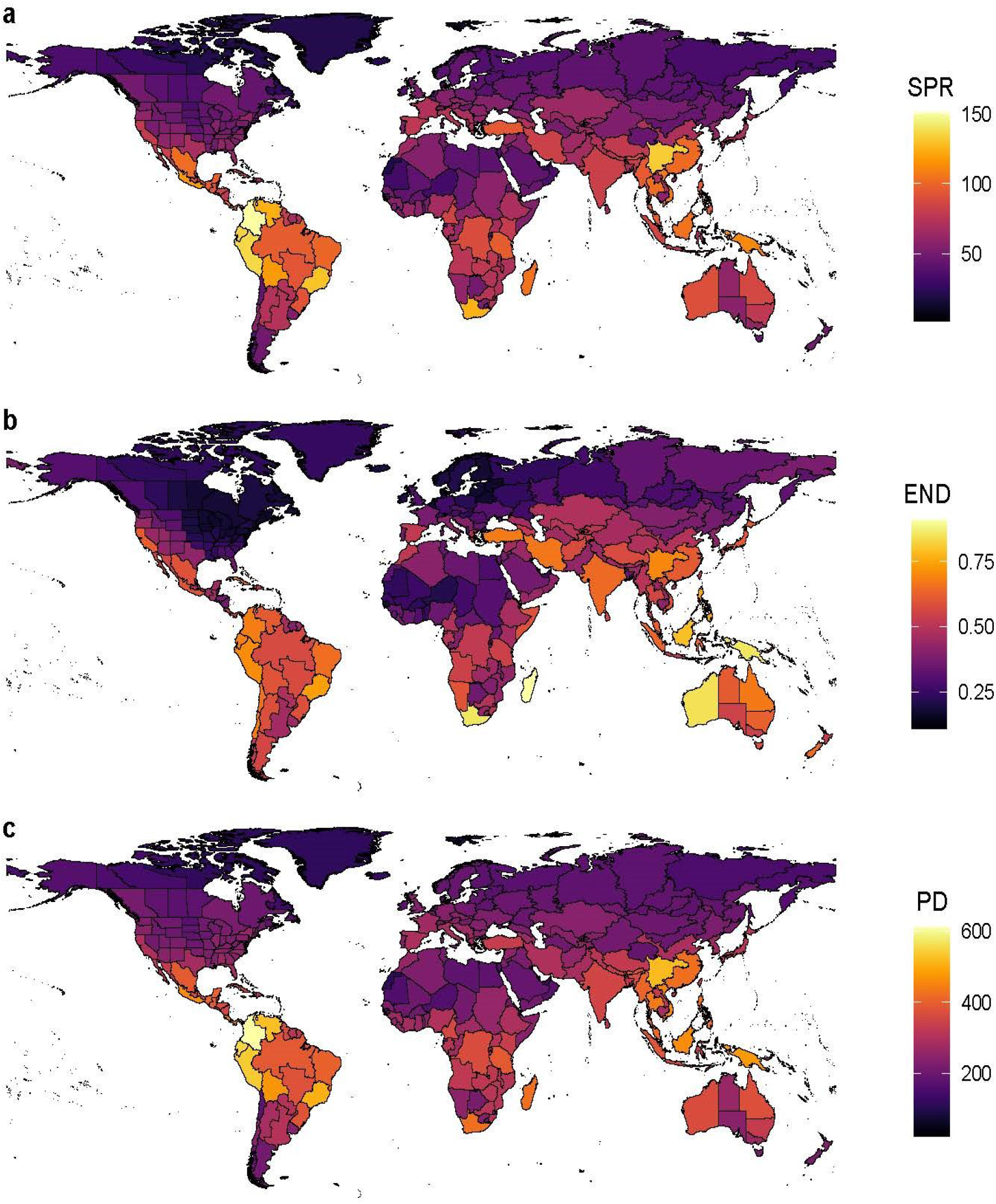
Current distribution of facets of plant diversity (SPR, END, PD) |. (**a**) polygons with the highest number of species (SPR). (**b**) polygons with the highest proportion of endemics (END) compared to the total number of species in that polygon. (**c**) polygons with the highest phylogenetic diversity (PD).

**Extended Data Figure 2.**
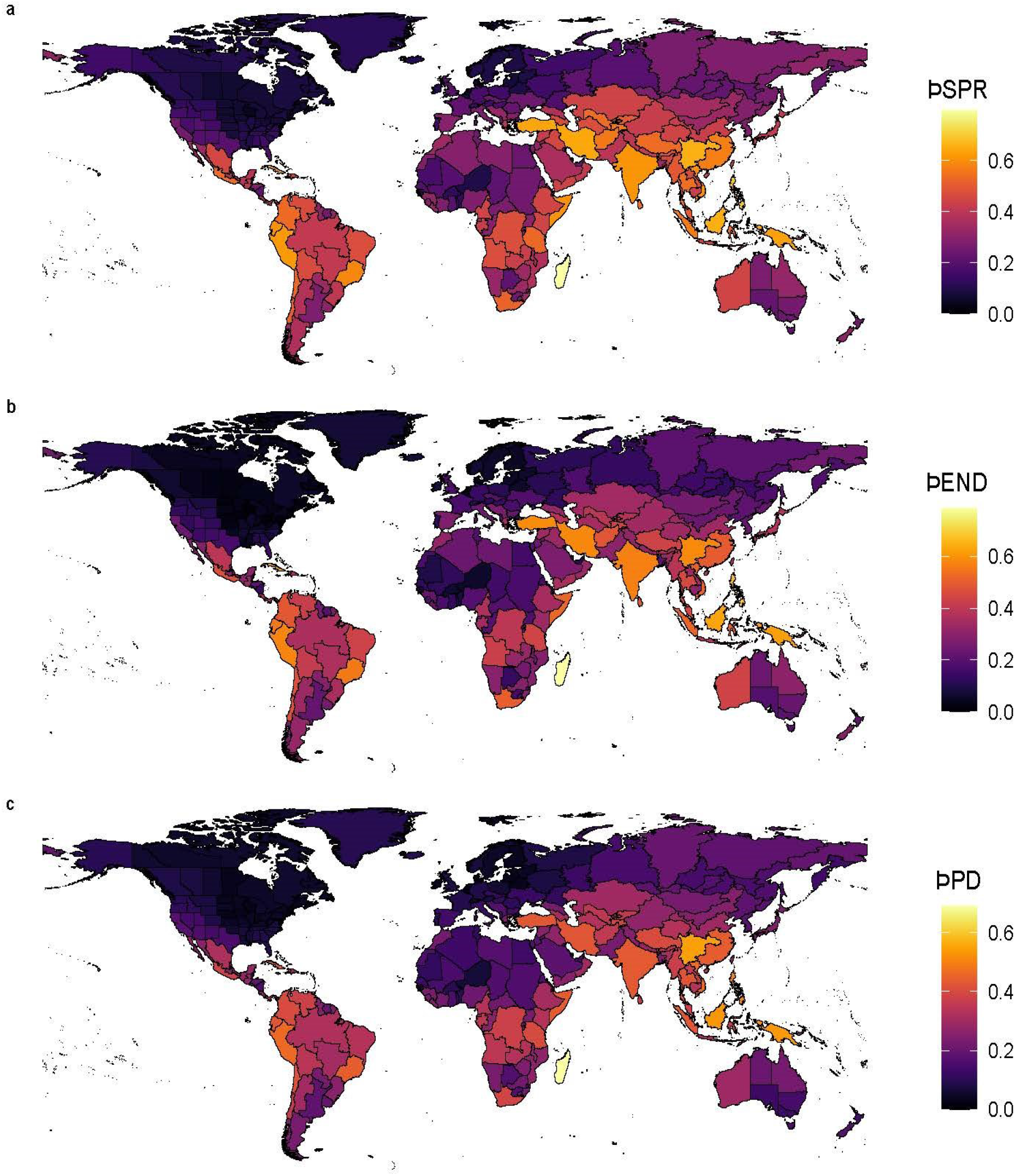
Distribution of plant diversity (ÞSPR, ÞEND, ÞPD) that is under threat |. (**a**) polygons with the highest proportion of threatened species (ÞSPR). (**b**) polygons with the highest proportion of threatened endemics (ÞEND) compared to the total number of species in that polygon. (**c**) polygons with the highest proportion of threatened phylogenetic diversity (ÞPD).

**Extended Data Figure 3.**
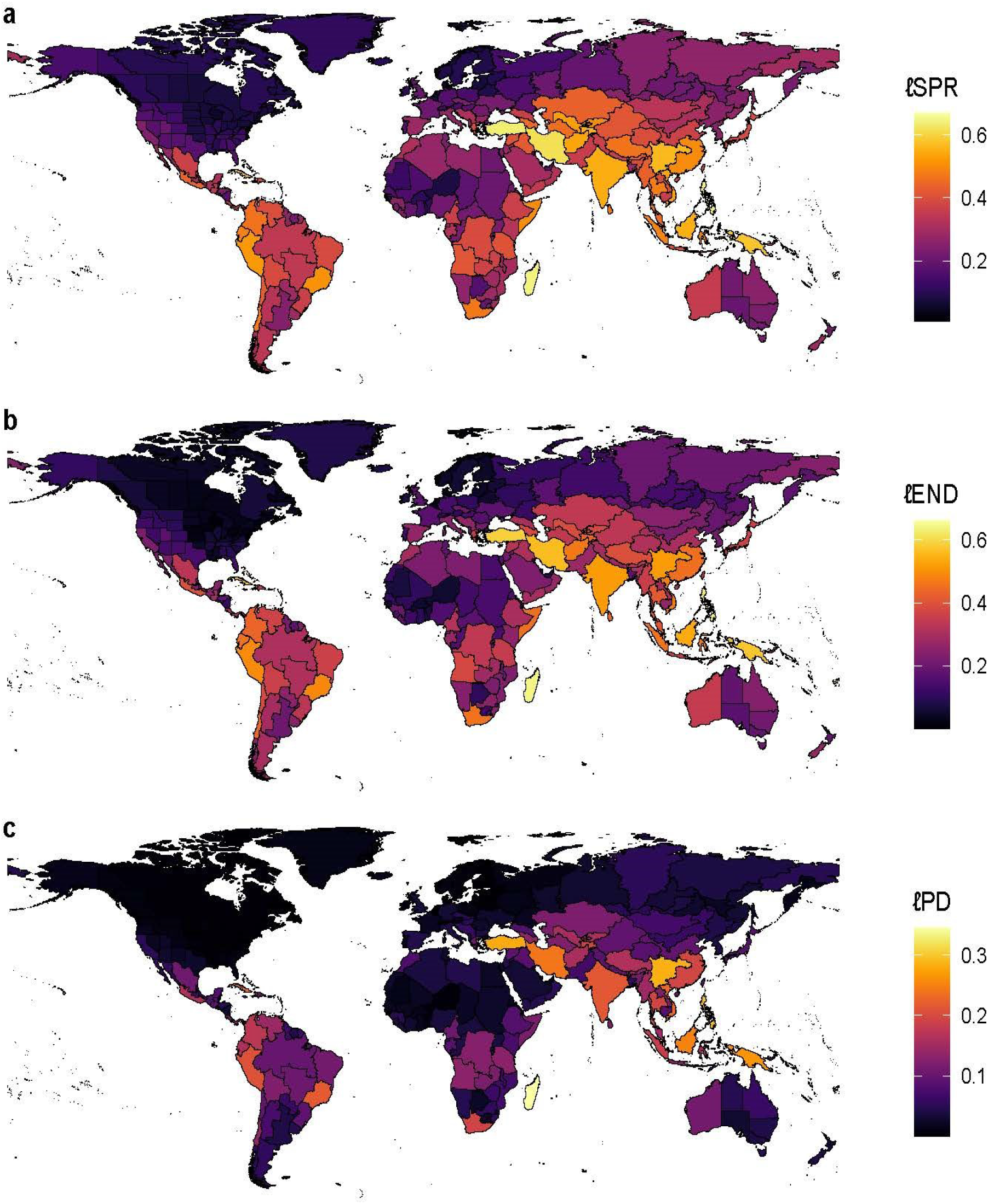
Expected biodiversity loss 50 years from now for each facet of plant diversity (ℓSPR, ℓEND, ℓPD) |. (**a**) polygons with the highest proportion of expected loss of species richness (ℓSPR). (**b**) polygons with the highest proportion of endemics expected to be lost (ℓEND) compared to its known flora. (**c**) polygons with the highest proportion of expected phylogenetic diversity loss (ℓPD).

**Extended Data Figure 4.**
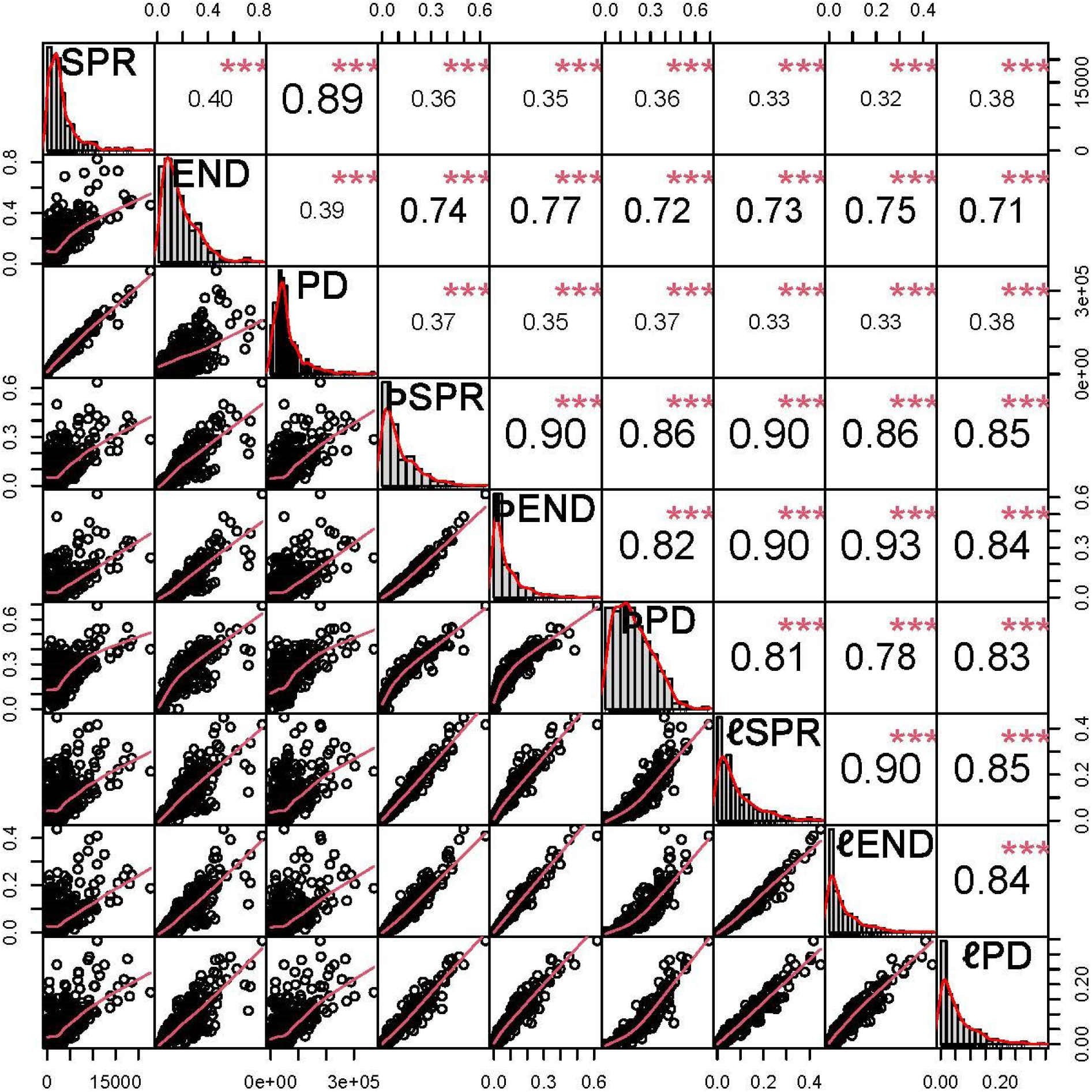
Correlation graphs for all variables. Current (SPR, END, PD), threatened (ÞSPR, ÞEND, ÞPD) and expected losses (ℓSPR, ℓEND, ℓPD), using Kendall’s τ, matching the correlogram presented in Fig. 1. (significance *** = 0.001, ** = 0.01, * = 0.05,. = 0.1,’’ = 1).

**Extended Data Figure 5.**
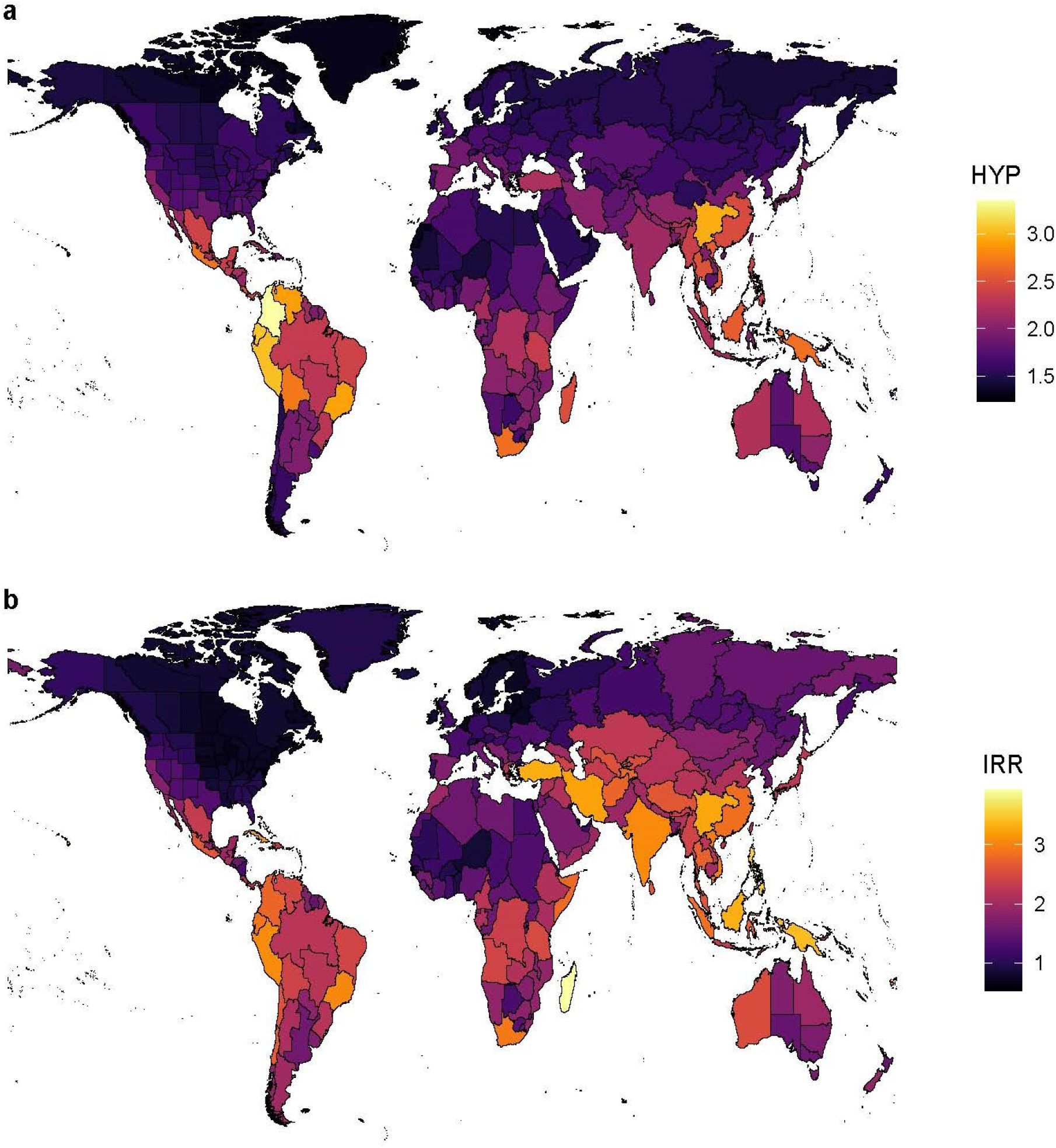
Top 2,5% and 5% hyperdiverse (HYP) and irreplaceable (IRR) areas, identified from groupings of biodiversity facets (see correlogram; Fig. 1)). (**a**) areas ranked by level of hyperdiversity, using values from axis 1 of the PCA analysis of PD+SPR. (**b**) areas ranked by level of irreplaceability and threat, using values from axis 1 of the PCA analysis of the other variables (ÞSPR, ℓSPR, END, ÞEND, ℓEND, ÞPD, ℓPD).

**Extended Table 1.**
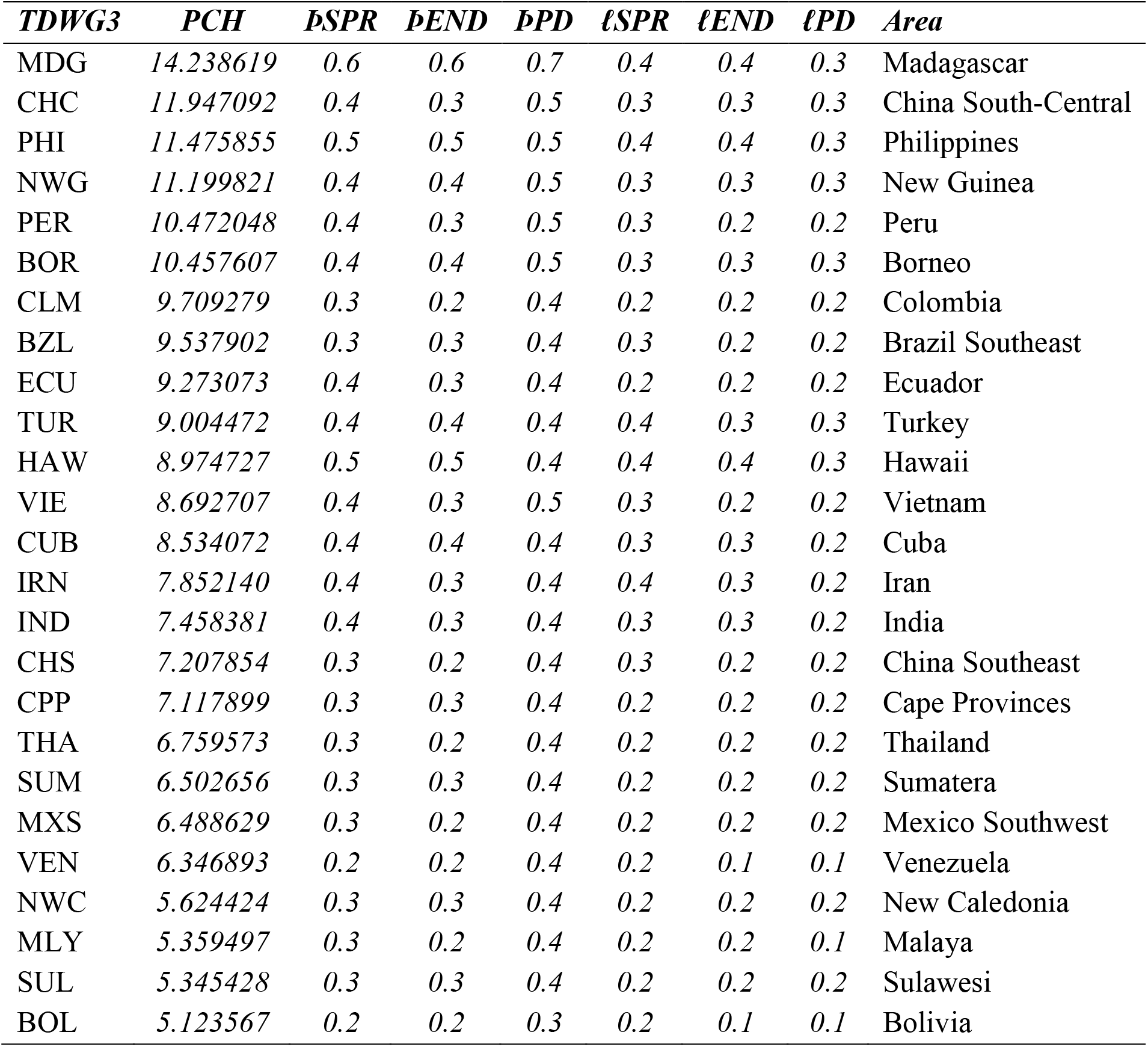
Top 25 PCH expressed by area. ‘TDWG3’ = TDWG level 3 area code, ‘Area’ = TDWG level 3 botanical country name (full ranking in Supplementary Table 3).

**Extended Data Table 2.**
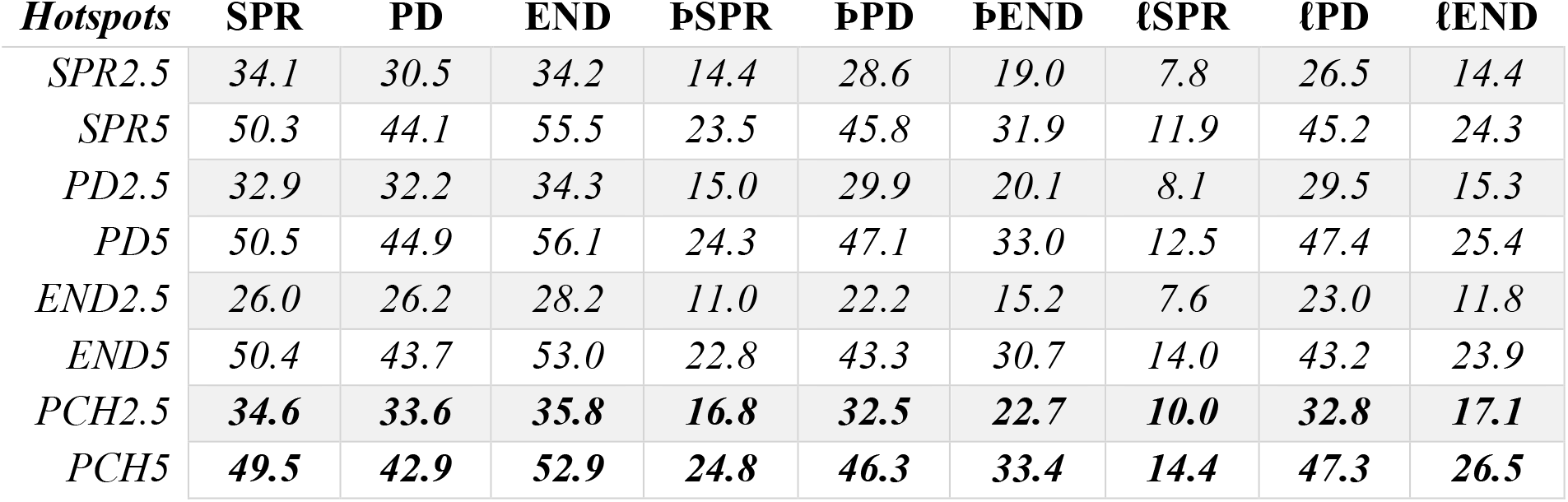
Respective contribution (percentage) of each biodiversity facet. Total contribution to current diversity measures (SPR, END, PD), threatened diversity (ÞSPR, ÞPD, ÞEND), and expected losses in 50 years time (ℓSPR, ℓPD, ℓEND) of the top 2.5% and 5% hotspots for each facet (SPR, PD, END), and for the top 2.5% and 5% PCH (calculations described in the Methods).

| *Hotspots* | **SPR** | **PD** | **END** | **ÞSPR** | **ÞPD** | **ÞEND** | **ℓSPR** | **ℓPD** | **ℓEND** |
| --- | --- | --- | --- | --- | --- | --- | --- | --- | --- |
| *SPR2.5* | 34.1 | 30.5 | 34.2 | 14.4 | 28.6 | 19.0 | 7.8 | 26.5 | 14.4 |
| *SPR5* | 50.3 | 44.1 | 55.5 | 23.5 | 45.8 | 31.9 | 11.9 | 45.2 | 24.3 |
| *PD2.5* | 32.9 | 32.2 | 34.3 | 15.0 | 29.9 | 20.1 | 8.1 | 29.5 | 15.3 |
| *PD5* | 50.5 | 44.9 | 56.1 | 24.3 | 47.1 | 33.0 | 12.5 | 47.4 | 25.4 |
| *END2.5* | 26.0 | 26.2 | 28.2 | 11.0 | 22.2 | 15.2 | 7.6 | 23.0 | 11.8 |
| *END5* | 50.4 | 43.7 | 53.0 | 22.8 | 43.3 | 30.7 | 14.0 | 43.2 | 23.9 |
| *PCH2.5* | **34.6** | **33.6** | **35.8** | **16.8** | **32.5** | **22.7** | **10.0** | **32.8** | **17.1** |
| *PCH5* | **49.5** | **42.9** | **52.9** | **24.8** | **46.3** | **33.4** | **14.4** | **47.3** | **26.5** |

**Extended Data Table 3.** Percentage of species in common. Percentage of species in common between each of the top 2.5% and 5% hotspots for each facet (SPR, END and PD) and PCH. ‘ANIDS’ = *n* species included (calculations described in the Methods).

| *Hotspots* | **SPR2.5** | **SPR5** | **PD2.5** | **PD5** | **END2.5** | **END5** | **PCH2.5** | **PCH5** | **ANIDS** |
| --- | --- | --- | --- | --- | --- | --- | --- | --- | --- |
| *SPR2.5* | 100.0 | 67.8 | 80.1 | 67.6 | 41.8 | 55.6 | 67.5 | 56.5 | 113141 |
| *SPR5* | 100.0 | 100.0 | 100.0 | 96.9 | 79.1 | 75.0 | 95.5 | 83.3 | 166852 |
| *PD2.5* | 77.2 | 65.3 | 100.0 | 65.1 | 49.0 | 59.2 | 86.1 | 60.2 | 109005 |
| *PD5* | 100.0 | 97.2 | 100.0 | 100.0 | 85.1 | 78.1 | 100.0 | 86.4 | 167391 |
| *END2.5* | 31.9 | 40.9 | 38.8 | 43.8 | 100.0 | 51.6 | 50.8 | 45.5 | 86245 |
| *END5* | 82.1 | 75.1 | 90.7 | 77.9 | 100.0 | 100.0 | 100.0 | 90.4 | 167005 |
| *PCH2.5* | 68.3 | 65.6 | 90.5 | 68.4 | 67.4 | 68.6 | 100.0 | 69.8 | 114577 |
| *PCH5* | 82.0 | 82.0 | 90.7 | 84.7 | 86.6 | 88.8 | 100.0 | 100.0 | 164142 |

**Extended Data Table 4.**
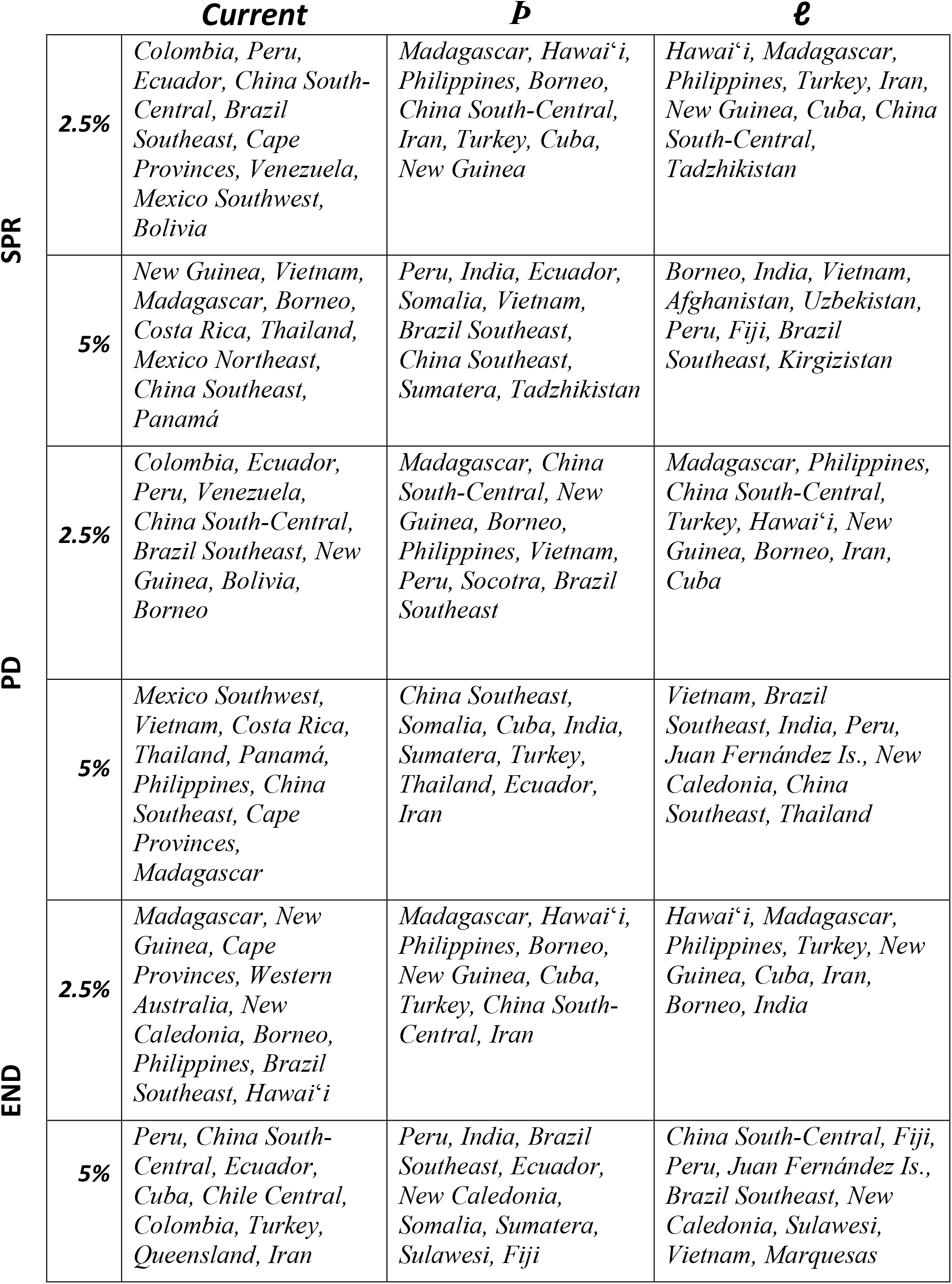
Hotspots for each facet. Botanical countries are ordered by decreasing rank (see **Supplementary Table 3** for values and full ranking).

